# A Casz1 - NuRD complex regulates temporal identity transitions in neural progenitors

**DOI:** 10.1101/2020.02.11.944470

**Authors:** Pierre Mattar, Christine Jolicoeur, Sujay Shah, Michel Cayouette

## Abstract

Neural progenitor cells alter their output over developmental time to generate different types of neurons and glia in the correct sequences and proportions. A number of ‘temporal identity factors’ that control transitions in progenitor competence have been identified, but the molecular mechanisms underlying their function remain unclear. Here, we asked how the transcription factor *Casz1*, the mammalian orthologue of *Drosophila castor,* regulates competence during retinal neurogenesis. We show that *Casz1* is required to control the transition between neurogenesis and gliogenesis. Using BioID proteomics, we reveal that Casz1 interacts with the nucleosome remodeling and deacetylase (NuRD) complex in retinal cells. Finally, we show that both the NuRD and the polycomb repressor complexes are required for Casz1 to promote the rod fate and suppress gliogenesis. As other temporal identity factors have been found to interact with the NuRD complex in other contexts, we propose that these factors might act through a common biochemical process to regulate neurogenesis.

## Introduction

During neurogenesis, virtually all neural stem and progenitor cells change their output dynamically over developmental time, first generating neurons, and then switching to generate glia [1]. In regions of the central nervous system (CNS) such as the neocortex and retina, progenitors undergo additional temporal identity transitions in competence to generate specific neuron subtypes at precise stages of development. In the vertebrate retina, retinal progenitor cells (RPCs) have two distinctive phases of multipotency [2–5]. Within each phase, RPCs can simultaneously produce a number of different neuronal and glial subtypes. For example, during embryonic stages, RPCs generate retinal ganglion cells, cone photoreceptors, horizontals, and amacrine neurons. As the generation of these cell types peaks, the production of rod photoreceptors increases, peaking at around birth. Postnatally, RPCs lose the competence to generate the early-born neuronal subtypes, and instead begin to produce bipolar neurons and finally Müller glia [2,4–8].

Importantly, while many fate determinants have been identified that are necessary for the generation of specific retinal cell types, many of these factors drive cell cycle exit, and are not compatible with regulating progenitor competence state [9–12]. It thus remains largely unclear how multipotent progenitors select between alternative differentiation programmes available to them at any given stage to generate lineages with correct cell numbers and birth order.

While our understanding of the molecular mechanisms that control temporal identity in vertebrate neural progenitors is limited [12], several transcription factor cascades controlling this process in *Drosophila* have been identified [13]. In the fly ventral nerve cord, landmark work has demonstrated that most neural stem cells express a sequence of transcription factors as development proceeds, namely: hunchback, Krüppel, nub/pdm2 (collectively pdm), castor, and grainyhead [13]. This cascade of ‘temporal identity factors’ acts as a general timing mechanism to coordinate the output of hundreds of individual neuroblast lineages within the *Drosophila* CNS. How this transcription factor cascade is able to integrate coherently into so many different lineages remains unclear at the mechanistic level.

To begin to identify molecular pathways involved in temporal transitions during neural development, we chose to focus on the zinc finger transcription factor *Casz1*, which is the murine orthologue of *Drosophila castor*. In fly neuroblasts, *castor* is amongst the most influential of the temporal factors [14], and we have previously reported that *Casz1* plays a conserved role in regulating progenitor potential in the developing mouse retina [15]. We and others have previously shown that Casz1 interacts physically and functionally with key subunits of the polycomb repressive complex (PRC) [16, 17]. Polycomb, in turn, is a well-known regulator of developmental timing in neurogenesis [18–21]. However, whether the Casz1-PRC interaction is functionally important in neural progenitors has not been addressed. Here, to identify the molecular pathway utilized by Casz1 to regulate neural progenitor competence, we used BioID proteomics to identify the retinal Casz1 interactome. We found that Casz1 associates primarily with the NuRD complex in retinal cells. Moreover, functional dissection of the Casz1 interactome demonstrates that both the NuRD complex and PRC are required for Casz1 to promote the rod photoreceptor cell fate and suppress the production of Müller glia. Thus, our data demonstrate functional interdependency between temporal identity factors and epigenetic regulators, suggesting a unifying model of how these two well-known temporal patterning systems operate.

## Results and Discussion

### Casz1 suppresses Müller glia production in postnatal retinal progenitors

We have previously shown that *Casz1* regulates the mid/late temporal identity in RPCs. During retinal development, the expression of *Casz1* mRNA and protein steadily increases in RPCs, peaks at around P0, and declines postnatally (Fig 1A) [3, 15]. We previously generated a *Casz1* conditional knockout (cKO) mouse, and found that *Casz1* is required in the retina to suppress the production of early-born neurons and late-born Müller glia, while promoting the production of rod photoreceptors [15]. As these previous experiments inactivated *Casz1* in early-stage RPCs, the specific requirement for *Casz1* in perinatal RPCs, which generate glial cells, remains unclear. To address this question, we introduced control (GFP) or Cre recombinase acutely into cohorts of P0 *Casz1^Flox/+^* or *Casz1^Flox/Flox^* RPCs using electroporation. We then cultured the resultant retinas *ex vivo* for 14 days, when retinal development is complete. No significant difference in cell production was observed in GFP transfections from either genotype. However, when Cre was transfected, rod photoreceptor production was significantly reduced in *Casz1^Flox/+^* retinas, and further reduced in *Casz1^Flox/Flox^* explants (Fig. 1B-D). These reductions were accompanied by concomitant increases in Müller glia production, which were ∼4-fold more abundant in *Casz1^Flox/Flox^* transfections. Consistently, RNA-Seq analysis of sorted RPCs at P2 revealed that expression of rod marker genes was reduced, whereas that of Müller marker genes was increased in *Casz1* cKOs compared to controls (Fig. 1E). We conclude that *Casz1* is required in postnatal RPCs to promote rods and suppress Müller glia production.

**Fig. 1.**
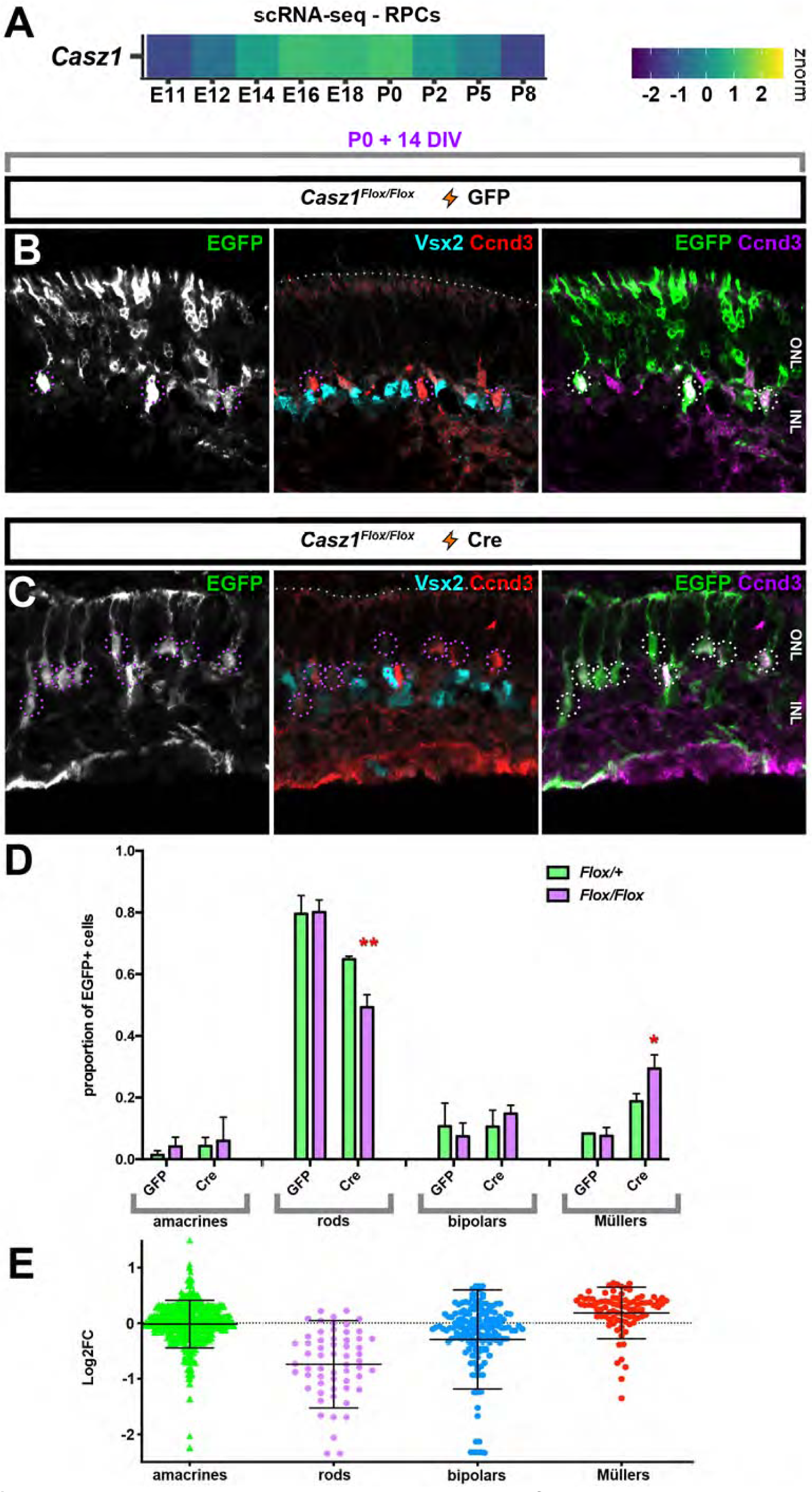
*Casz1* controls the rod versus Müller glia fate decision in postnatal retinal progenitors. (A) Relative *Casz1* mRNA expression levels in RPCs during development [3]. (B, C) P0 *Casz1^Flox/Flox^* or *Casz1^Flox/+^* retinal progenitors were transfected with control (B) or Cre (C) constructs, and cultured *ex vivo* for 2 weeks to allow development to reach completion. Explants were then harvested, sectioned, and stained for Vsx2 and Ccnd3 proteins, which mark Müller glia and bipolar cells. (D) Quantitation of the cell-type composition of the resultant transfections. See Table S1 for statistical summary. * p < 0.05; ** p < 0.01. (E) Re-analysis of previously published RNA-seq data from sorted P2 *Casz1* cKO vs. control RPCs[16]. Plot depicts Log_2_ fold-change in gene expression for cell subtype marker genes based on previously published cell-type profiling [59, 60]. ONL, outer nuclear layer; INL, inner nuclear layer.

### Casz1 interacts with the NuRD complex in RPCs

Having timed a requirement for *Casz1* to perinatal stages of development, we reasoned that we could take advantage of the quantitatively large number of RPCs generating rods and Müllers at these stages to identify Casz1 co-factors via biochemical purification. Unfortunately, we found that despite robust mRNA expression levels, Casz1 protein was not detectable in western blots via standard methods using several different antibodies. We only observed full-length Casz1 isoforms when protein lysates were stabilized by formaldehyde cross-linking [16]. Since we could not purify native Casz1-containing protein complexes, we turned to BioID proteomics. The BioID approach bypasses the requirement to purify target proteins directly. Instead, the target protein is tagged with a BirA biotin ligase domain (BirA*) containing a mutation that abolishes target selectivity. BioID relies on the preferential biotinylation of the interactome as a function of molecular proximity. BirA* was fused to the N-terminus of Casz1v2 (Fig. 2A, B). We focused on Casz1v2 since our previous experiments indicated that it was the splice variant involved in rod production [16].

**Fig. 2.**
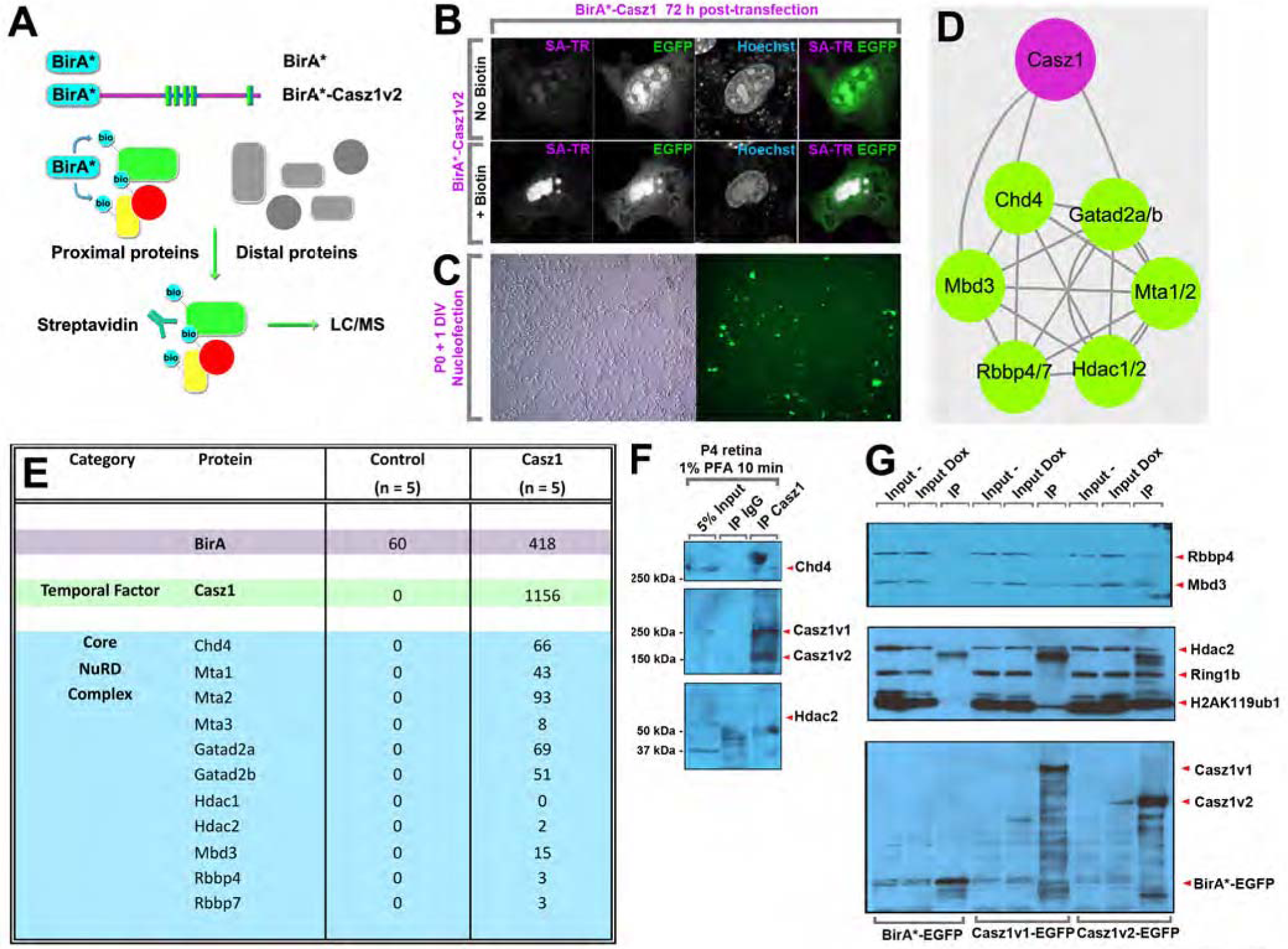
BioID proteomics identifies Casz1 interacting proteins from cultured retinal progenitors. (A) BioID strategy. Proteins are tagged with a mutant version of the bacterial biotin ligase BirA (BirAR118G, also called BirA*). BirA* promiscuously biotinylates all proteins with a free primary amine group (ie. lysine residues). Biotinylated proteins are isolated using streptavidin coupled beads and subjected to liquid chromatography, tandem mass spectrometry (LC/MS-MS). (B) Representative images of the nuclear localization of BirA*-tagged Casz1v2 constructs transfected into 3T3 fibroblasts and cultured with or without exogenous biotin. Biotinylated proteins are visualized using Texas Red-conjugated streptavidin (SA-TR). (C) Transfection of BioID constructs into dissociated retinal progenitor cultures using nucleofection. (D) Model of Casz1 interactome. (E) Summary of pep ide counts. (F) Validation of Casz1/NuRD complex interactions using cross-linking assisted immunoprecipitation from P4 retinas. (G) Validation of Casz1/NuRD complex interactions using stably expressing 293Trex cell lines. EGFP-fused control or Casz1 protein complexes were immunoprecipitated using anti-GFP antibodies.

Next, we introduced nuclear-localized control BirA* [22] or Casz1v2-BirA* constructs into primary retinal cultures by nucleofection. Transfected retinal cells were cultured at high-density using a defined serum-free medium [6, 23] (Fig. 2C). Cultures were supplemented with exogenous biotin for 6 h and harvested. Since Casz1 is a nuclear protein in the retina[15, 16], we additionally extracted nuclei, as we observed considerable amounts of cytoplasmic biotin when untransfected control cells were stained with fluorochrome-conjugated streptavidin (Fig. 2B). Casz1 had previously been observed to interact with components of PRC1, both in cultured cells and in adult retinas [16, 17]. To our surprise, when we examined the BioID interactome, we saw little evidence for association between Casz1 and the PRC in perinatal retina. Polycomb-associated proteins such as Cbx1/3/5, Atrx, and Hmgb1/2/3 were detected exclusively in the Casz1 interactome, but with only a few peptides each. Moreover, core subunits such as Ring1, Rnf2, Pcgf and Phc proteins were not detected. Instead, the NuRD complex was the most prominent interacting partner identified. All of the core subunits of the complex were observed, including Chd4, Mbd3, Gatad2a/b, and Mta1/2. Subunits such as Rbbp4/7 and Hdac2 were also observed, albeit with only a few peptide spectra in total. None of these proteins were ever observed in BioID complexes purified from control transfections (Fig. 2D, E).

To validate the observed interactions, we performed several experiments. To examine the association between Casz1 and the NuRD complex in retinal tissue, we first performed cross-linking assisted immunoprecipitation using the RIME proteomic workflow [24], which we have previously reported stabilizes Casz1 protein complexes [16]. As expected, Casz1 protein complexes purified from developing retinas contained both Chd4 and Hdac2 (Fig. 2F). Next, we performed conventional co-immunoprecipitation experiments using 293Trex cell lines that were engineered to stably and inducibly express control, Casz1v1-, or Casz1v2-EGFP fusion proteins. Immunoprecipitation with anti-GFP antibodies revealed that NuRD complex members like Hdac2, Mbd3, and Rbbp4 were associated with both Casz1v1 and Casz1v2 isoforms (Fig. 2G). In contrast, the core polycomb subunit Rnf2 and the polycomb mark H2AK119ub1 were enriched by Casz1v2 but not Casz1v1, as we previously reported (Fig. 2G) [16].

Next, we assessed the co-localization of Casz1, Chd4, and Hdac proteins in retinal progenitors using immunohistochemistry. We observed that Casz1 was expressed throughout the nucleus, but accumulated on the outer rims of chromocenters, which are focal accumulations of pericentromeric, constitutive heterochromatin (Fig. S1A). This staining pattern disappeared in *Casz1* cKO RPCs (Fig. S1A), confirming specificity of the signal. The outer rims of chromocenters are also enriched for heterochromatic markers, such as Cbx1 and H3K9me3, which were accordingly detected in the Casz1 interactome (Fig. S1B, C). When we performed immunohistochemistry for partner proteins such as Chd4 or Hdac2, we observed that both proteins co-localized with Casz1 in nuclear foci that were usually in apposition with chromocenters (Fig. S1D, E). This suggests that Casz1 is localized to at least a subset of “NuRD bodies” [25] in RPCs.

To determine whether Casz1 regulates the localisation of NuRD proteins, we overexpressed Casz1v2 in RPCs using electroporation in retinal explants, and examined the distribution of Chd4 and Hdac2. We have previously found that Casz1v2 binds and expands chromocenters when overexpressed [16]. Accordingly, endogenous Chd4 and Hdac2 were similarly accumulated upon Casz1 transfection (Fig. S2). These data suggest that Casz1 can relocalize the NuRD complex within the nucleus.

Consistent with these results, temporal transcription factors have been previously been observed to associate with NuRD and PRC in other contexts. Indeed, *Drosophila* dMi-2 was originally discovered in a screen for hunchback interacting proteins, and was shown to function in the polycomb pathway to repress *Hox* genes [26]. Similarly, in lymphocyte development, both NuRD and PRCs are associated with Ikzf1 [27–29], which was shown to confer early temporal identity in the developing mouse retina [30]. Casz1 was also shown to interact with both NuRD and PRC in cultured cell lines [16,17,31], and temporal factors such as *Drosophila* grainyhead have been found interacting with the PRC [32, 33]. Similarly to mammalian neurogenesis, PRCs have also been shown to regulate competence transitions in *Drosophila* [34]. These observations suggest that temporal transcription factors might use a common biochemical pathway to regulate competence transitions during neural development, but functional data to explore this possibility is still missing.

### Casz1 requires the NuRD complex to promote rods and suppress gliogenesis

We therefore next set out to determine the functional requirement for the NuRD complex in Casz1-mediated neural cell fate decisions. Interestingly, while the NuRD complex had not previously been implicated in retinal neurogenesis, Hdac proteins were previously shown to be required for retinal progenitor proliferation and rod photoreceptor gene expression [35–37]. However, rod photoreceptor transcription factors that might partner with Hdacs to achieve these effects have not been described. To address the requirement for histone deacetylases for Casz1 function, we introduced control or Casz1v2 –expressing retroviruses into P0 retinal progenitors. After 4 h, we added vehicle, 50 nM trichostatin A (TSA) – a broad spectrum histone deacetylase inhibitor, or 500 nM UF010, a selective inhibitor of class I Hdacs - to the cultures for 4 days (Fig. 3A). Next, the drugs were washed out, and retinas were cultured *ex vivo* for 10 additional days to allow neurogenesis to reach completion. We then fixed the tissue explants and reconstructed the clones (Fig. 3B-G). Similar to our previous results with embryonic clones [15], and conversely to what we observed by conditional ablation of *Casz1* at P0 (Fig.1), Casz1v2 overexpression at P0 significantly increased the proportion of rods produced at the expense of Müller glia (Fig. 3H, I; Table S2). 50 nM TSA had no significant effect on rod vs. Müller production in control clones, but UF010 led to a significant reduction in rod production with a concomitant increase in Müllers. When Casz1-transduced clones were treated with TSA or UF010, Casz1v2-mediated rod production was completely lost, and Müller production increased significantly (Fig. 3H, I; Table S2).

**Fig. 3.**
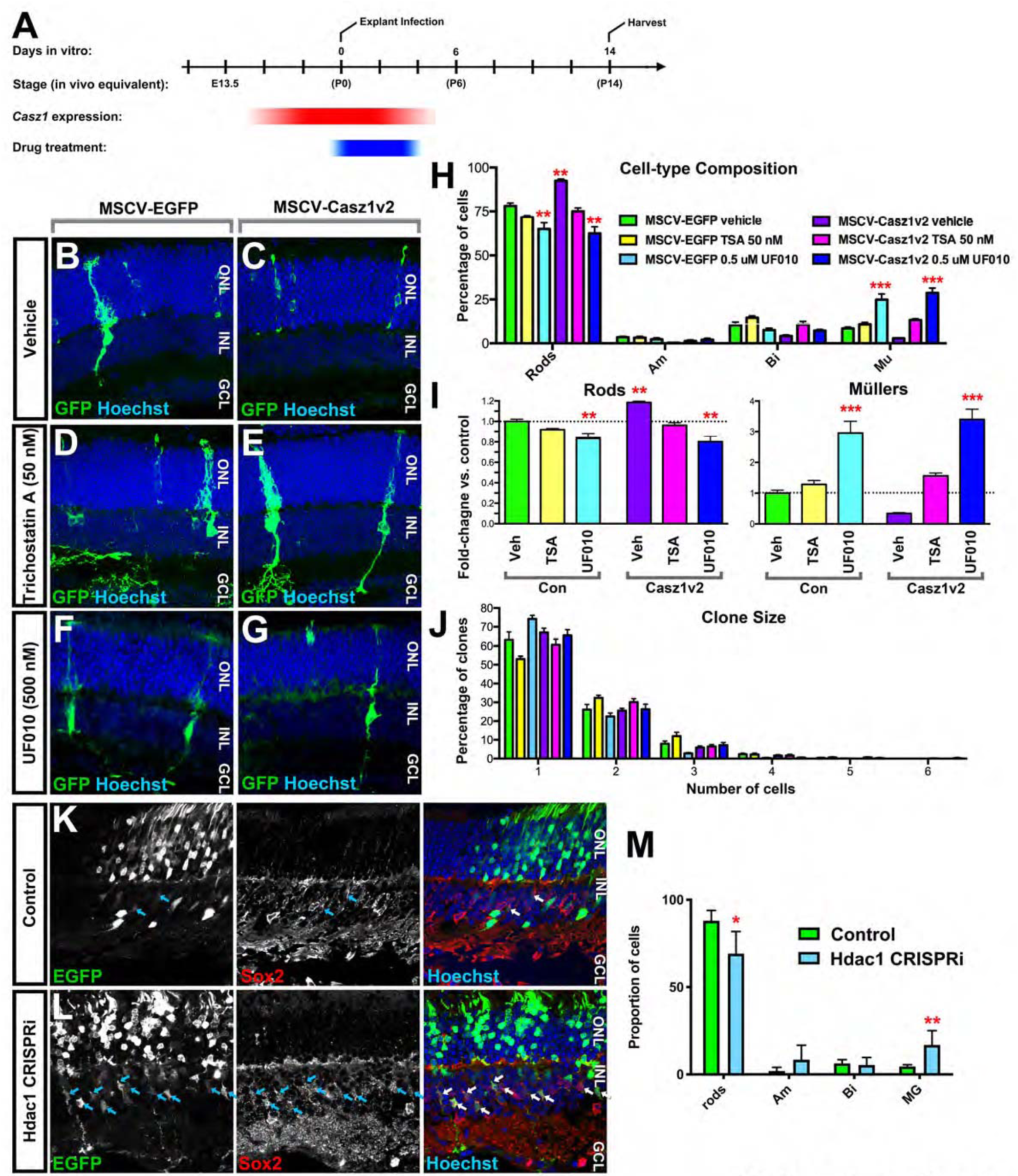
Casz1 requires histone deacetylase activity to control RPC output. (A) Experimental outline. Retroviruses were introduced at P0 at clonal density. Hdac inhibitors were added after 2-4 hours, and left in place for an additional 4 days. Explants were harvested and analyzed after 14 days *in vitro*. P0 RPCs were transduced with vector control (MSCV-EGFP) or Casz1v2 –expressing retroviruses in the presence or absence of 50 nM trichostatin A for 4 days. Retinas were cultured *ex vivo* for an additional 10 days to allow neurogenesis to reach completion. (B-G) Examples of control or Casz1v2 transduced retroviral clones cultured in the presence of vehicle (B, C), 50 nM TSA (D, E), or 500 nM UF010 (F, G). Infected cells were stained with antibodies against GFP (green) and DNA was stained with Hoechst 33342 (blue). (H, I) Cell-type composition of retroviral clones. See Table S2 for statistical summary. ** p < 0.01; *** p < 0.001. (J) Distribution of cells per clone. (K-M) Hdac1 CRISPRi via retinal electroporation at P0. Retinas were harvested at P21 and stained for Sox2 and Hoechst. (K) Control or (L) Hdac1 CRISPRi. (M) Quantitation of transfected cell fate. See Table S3 for statistical summary.

To determine which class I Hdac proteins were required for Casz1 functions, we focused on Hdac1. We used retinal electroporation to perform CRISPR interference (CRISPRi) on Hdac1 *in vivo*. Retinas were co-transfected at P0 with constructs encoding GFP, dCas9-KRAB-MeCP2 [38], and a guide construct targeting the transcriptional start of Hdac1. Retinas were harvested at P21, and the proportions of different transfected cell types were enumerated. We found that, similarly to pharmacological inhibition, Hdac1 CRISPRi significantly reduced the proportion of rods produced by Casz1v2 expression, while Müller glia were significantly increased (Fig. 3K-M, Table S3). Together, these data indicate that Casz1 requires histone deacetylase activity to suppress gliogenesis and promote rod production.

### Casz1v2 requires the polycomb repressive complex to promote rods and suppress gliogenesis

Next, we addressed the requirement for PRC activity in Casz1-dependent fate decisions. Our previous functional work suggested that Casz1v2 requires the PRC to safeguard transcription in rod photoreceptors, and previous biochemical work showed that Casz1 can associate with PRC (Fig. 2F) [16, 17]. Using BioID, we found little evidence for a direct interaction between Casz1 and PRC components, but the NuRD complex is known to physically and functionally associate with PRC, and the PRC has also previously been shown to oppose gliogenesis [18,26,39–41], and regulate temporal patterning in neural development [19,20,34,42], suggesting a possible indirect role for PRC in Casz1-mediated functions.

To test whether the PRC is required for Casz1 function, we utilized retroviral shRNAs targeting Ring1 or Rnf2 [16], which are necessary to mediate PRC activity. These retroviruses were first introduced into retinal explants. After 14 days *in vitro*, explants were harvested, and clones were reconstructed (Fig. 4A, B). Similarly to *Casz1* loss-of-function, we found that shRNAs targeting Ring1 or Rnf2 promoted gliogenesis and reduced the production of rod photoreceptors (Fig. 4C; Table S4). Next, we performed combinatorial experiments, expressing Casz1v2 and shRNA combinations using electroporation. When Casz1v2 was co-expressed with Ring1 or Rnf2 shRNA constructs, Casz1v2-mediated suppression of Müller gliogenesis was reversed (Fig. 4E, F; Table S5). These were specific effects, as they could be rescued by co-expressing a human *Rnf2* cDNA that could not be knocked down due to mismatches with the mouse shRnf2-389 hairpin [16]. We conclude that Casz1v2 requires PRC downstream of the NuRD complex to mediate its effects on cell fate.

**Fig. 4.**
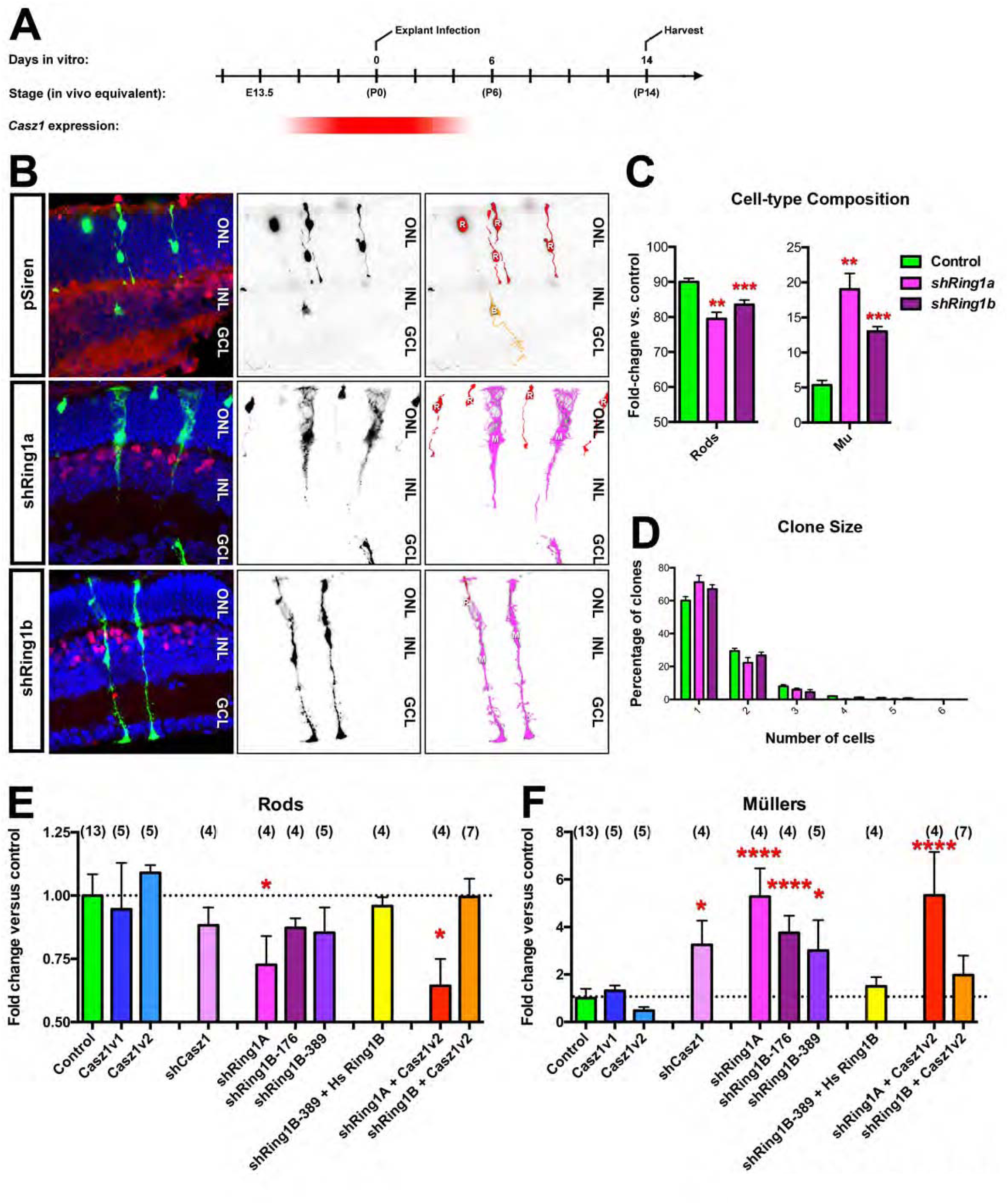
Casz1 requires polycomb to control retinal progenitor output. (A) Experimental timeline. (B) Examples of vector control (pSiren) or shRNA clones that were transduced at P0 and cultured for 14 days *in vitro*, marked by ZsGreen expression (green). Sections were stained for Vsx2 (red) and Hoechst (blue). (C, D) Cell-type composition (C) and clone size distribution (D) of the resultant clones (n=4). (E, F) Combinatorial expression/co-expression of Casz1 and polycomb loss or gain –of-function constructs. P0 retinas were electroporated with the indicated construct combinations, and harvested after 14 days *in vitro*. The proportion transfected rods (E) or Müller glia (F) was quantitated and normalized to the proportions obtained in control transfections. The control was pooled from overexpression (pCIG2: n=8) and shRNA (pSiren: n=5) vector transfections. The asterisks denote significant differences versus control as determined by one-way ANOVA with Tukey’s post-hoc test. n-values are presented in parentheses. * p < 0.05; ** p < 0.01; *** p < 0.001; **** p < 0.0001. (E) Rod production in Casz1v2 transfections was significantly different versus Casz1v2 + shRing1a (p < 0.0001). (F) Müller glia production in Casz1v2 transfections was significantly different versus Casz1v2 + shRing1a (p < 0.0001) and Casz1v2 + shRnf2-176 (p < 0.05).

Our data suggest a model where Casz1 interacts directly with the NuRD complex, which leads to subsequent PRC recruitment. NuRD and PRC have been previously shown to regulate neocortical development [18,20,43,44] and the PRC has been previously implicated in retinal development [39,45–47]. Physical and functional interactions between NuRD complexes and PRCs have been extensively described, albeit chiefly for PRC2 [40,48,49] whose core subunits are Ezh1/2, Suz12, Eed, and Rbbp4/7. Indeed, Rbbp4/7 proteins are also shared with the NuRD complex. Moreover, direct physical interactions between NuRD and PRC components have been previously observed in neural progenitors [41, 48]. These studies suggested that NuRD/PRC cooperate to oppose gliogenesis. Importantly, other studies have shown that the complexes are also required for progenitors to undergo competence transitions and can thereby abnormally prolong neurogenesis and oppose gliogenesis [18, 21]. Taken together, these experiments argue that NuRD and PRC cooperate to regulate developmental timing in the CNS, but that the specific activities of the complexes are context dependent. It will be interesting to examine whether these complexes play different roles during other stages of retinal development.

Casz1 has previously been shown to function as a critical regulator of cardiac development and blood vessel formation [50–52], but the molecular mechanisms involved remain unclear. It will be interesting to determine whether a similar Casz1-mediated recruitment of the NuRD complex is also at play in these contexts. Additionally, since Casz1 expression is maintained in mature photoreceptors, our data suggest that Casz1 might be important for organizing Hdacs to promote rod photoreceptor gene expression. Intriguingly, Casz1 is essential for rod photoreceptor viability [16], and Hdac proteins become dysregulated in some types of photoreceptor degeneration [53, 54], raising the possibility that Casz1 downregulation or dysfunction might contribute to these diseases.

It was recently proposed that cell fate decisions in *Drosophila* neuroblasts involve the interplay of spatial transcription factors and temporal transcription factors, where spatial factors would act on the epigenome to modify target gene access for the temporal factors. The temporal factors would then generate different cell fates in a step-wise manner [13]. Our data suggest a model with reversed logic: that temporal factor cascades utilize a common molecular pathway to perform step-wise modifications to the epigenome, which can then be read out by lineage-specific cell fate determinants – i.e. the spatial factors – to generate specific neuronal identities. We believe this model might explain i) how temporal transcription factor cascades can operate coherently in so many contexts, and ii) why their sequential expression in neural progenitors has been conserved in evolution.

## Methods

### Animal care

Mouse husbandry was performed in accordance with the guidelines of the uOttawa or IRCM animal care committee and the Canadian Council on Animal Care. CD1 mice were obtained from Taconic. *Casz1^Flox/Flox^* [15] and *R26-Stop-EYFP* [55] alleles and genotyping protocols were previously described. See also Table S6.

### Plasmids

Control and Casz1v2 retroviral (pMSCV-EGFP vector) and electroporation (pCIG2 vector) constructs were previously described [15]. shRNA and overexpression constructs for Rnf2 and Ring1 were previously described [16]. pCag-Cre was was a gift from Connie Cepko (Harvard University; Addgene plasmid # 13775). dCas9-KRAB-MeCP2 was a gift from Alejandro Chavez and George Church (Harvard University; Addgene plasmid # 110821). The guide RNA construct was generated by excising Cas9 from pX330-U6-Chimeric_BB-CBh-hSpCas9 via restriction digest, which was a gift from Feng Zhang (Addgene plasmid # 42230). pDEST-pcDNA5-FLAG-NLS-BirA was a gift from Ann-Claude Gingras (Lunenfeld-Tanenbaum Research Institute) [22]. BirA*-Casz1v2 was generated by PCR cloning.

### Transfection, transduction, and culture

*In vivo* and *ex vivo* retinal electroporation were performed as described previously [15, 16]. Retroviral preparation, *ex vivo* retroviral transduction, and clone reconstruction were performed as described previously [15]. Histone deacetylase inhibitors were administered 2-4 h after virus administration to ensure that viral integration was not altered. Trichostatin A was purchased from New England Biolabs (9950). UF010 was purchased from Cayman Chemical (21273). UF010 stocks were discarded after 2 weeks. In both cases, fresh drug was applied daily, and washed out after the 4^th^ day. 293Trex stable cell lines were generated as described [56], except that we expanded our cell lines from single cell clones. 3T3 cell transfection and microscopy were performed as described previously [16]. 1 µg/ml doxycycline (Alfa Aesar J60579) was added 24 h prior to harvest.

### Biochemistry

Crosslinking-assisted immunoprecipitation was performed according to the RIME workflow [24]. Protein immunoprecipitation was performed as described previously [16]. Antibodies are listed in Table S7.

### BioID Proteomics

One P0 mouse litter (∼20 eyes) was used for each experimental replicate. Retinas were dissected and dissociated as previously described [6], and nucleofected with BioID constructs using an Amaxa Nucleofector II and the Mouse Neural Stem Cell Nucleofector Kit (Lonza; VPG-1004) according the manufacturer’s instructions. Cells were plated on poly-L-lysine/laminin coated dishes in RGM medium [6, 23], and cultured for 48 h prior to harvest. The culture medium was supplemented with exogenous biotin (50 μM final concentration) 6 h prior to harvest.

Cells were washed 3x with PBS with 100 μM PMSF. Next, nuclei were harvested using cell lysis buffer (5 mM PIPES, pH 8.0, 85 mM KCL, 0.5% NP40, and Roche cOmplete, Mini, EDTA-free protease inhibitors). Cells were then scraped and collected, and the nuclei extracted for a further 15 minutes on ice with agitation. Lysis, sonication, bead coupling, and washing steps were performed as previously described [57], using 60 μl of MyOne Streptavidin Dynabeads (Invitrogen).

Proteins were digested on-bead with 0.5 ug Sequencing Grade Modified Trypsin (Promega) overnight at 37°C with agitation. The supernatants were collected and the beads were additionally washed two times with 100 μl ddH_2_0. Supernatants were pooled, reduced and alkylated with 9 mM dithiothreitol at 37°C for 30 min followed by 17 mM iodoacetamide at room temperature for 20 min in the dark. Supernatants were acidified with trifluoroacetic acid and residual detergents removed using a Waters Oasis MCX 96-well Elution Plate following the manufacturer’s instructions. Samples were eluted in 10% ammonium hydroxide/90% methanol (v/v), dried, and reconstituted under agitation for 15 min in 15 µL of 5% FA.

### LC-MS/MS analysis

LC was performed using an LTQ Orbitrap Velos (ThermoFisher Scientific) equipped with a Proxeon nanoelectrospray Flex ion source as described [56]. Briefly, we used PicoFrit fused silica capillary columns (15 cm x 75 µm i.d; New Objective, Woburn, MA) with C-18 reverse-phase material and high pressure packing cells (Jupiter 5 µm particles, 300 Å pore size; Phenomenex, Torrance, CA). LC-MS/MS data was acquired using a data-dependent top11 method combined with a dynamic exclusion window of 30 sec. Bioinformatic analysis was performed using Proteome Discoverer (version 2.1) and Mascot 2.5 (Matrix Science) against the Uniprot database. Data analysis was performed using Scaffold (version 4.5.3).

### Histology and Microscopy

Immunohistochemistry was performed as previously described [15, 58]. Microscopy was performed using Zeiss LSM700, LSM710 (IRCM), or LSM880 instruments (CBIA Core, uOttawa) using Zen (Zeiss), Volocity (Perkin Elmer), Fiji (ImageJ), and Adobe Photoshop (Adobe) software. Primary antibodies are listed in Table S7.

### RNA-seq analysis

RNA-seq on control and *Casz1* cKO RPCs was previously published (<u>GSE115778</u>). To examine cell-type-specific gene expression, we adapted scRNA-seq cluster data [59], selecting genes with “MyDiff” scores greater than 1.4. For rod photoreceptor genes, we supplemented the list with additional rod genes from a previous study [60].

### Quantitation and Statistical Analysis

Statistical analyses were performed using GraphPad Prism software. n-values refer to independent experiments and/or animals as described in the text.

## Acknowledgements

We thank Brian Clark and Seth Blackshaw and Johan Ericsson for sharing scRNA-seq data and antibodies. We thank Alejandro Chavez, Connie Cepko, George Church, Ann-Claude Gingras, and Feng Zhang for sharing plasmids. We thank Jean-François Côté for providing his BirA*-EGFP 293 tRex cell line. We thank Thanh Dang for technical assistance, and Jessica Barthe, Marie-Claude Lavallée for help with animal husbandry. We also thank Denis Faubert and Josée Champagne of the IRCM proteomic core facility for help with the BioID experiments, and members of the Cayouette and Mattar labs for insightful discussions. This work was supported by Fighting Blindness Canada and an operating grant from the Canadian Institutes of Health Research (CIHR, FDN 159936). P.M. was supported by a CIHR Postdoctoral Fellowship and currently holds the Gladys and Lorna J. Wood Chair for Research in Vision. M.C. holds the Gaëtane and Roland Pillenière Chair in Retina Biology and is a Research Scholar Emeritus of the Fonds de recherche du Québec – Santé.

## Supplementary Figures and Tables

**Fig. S1.**
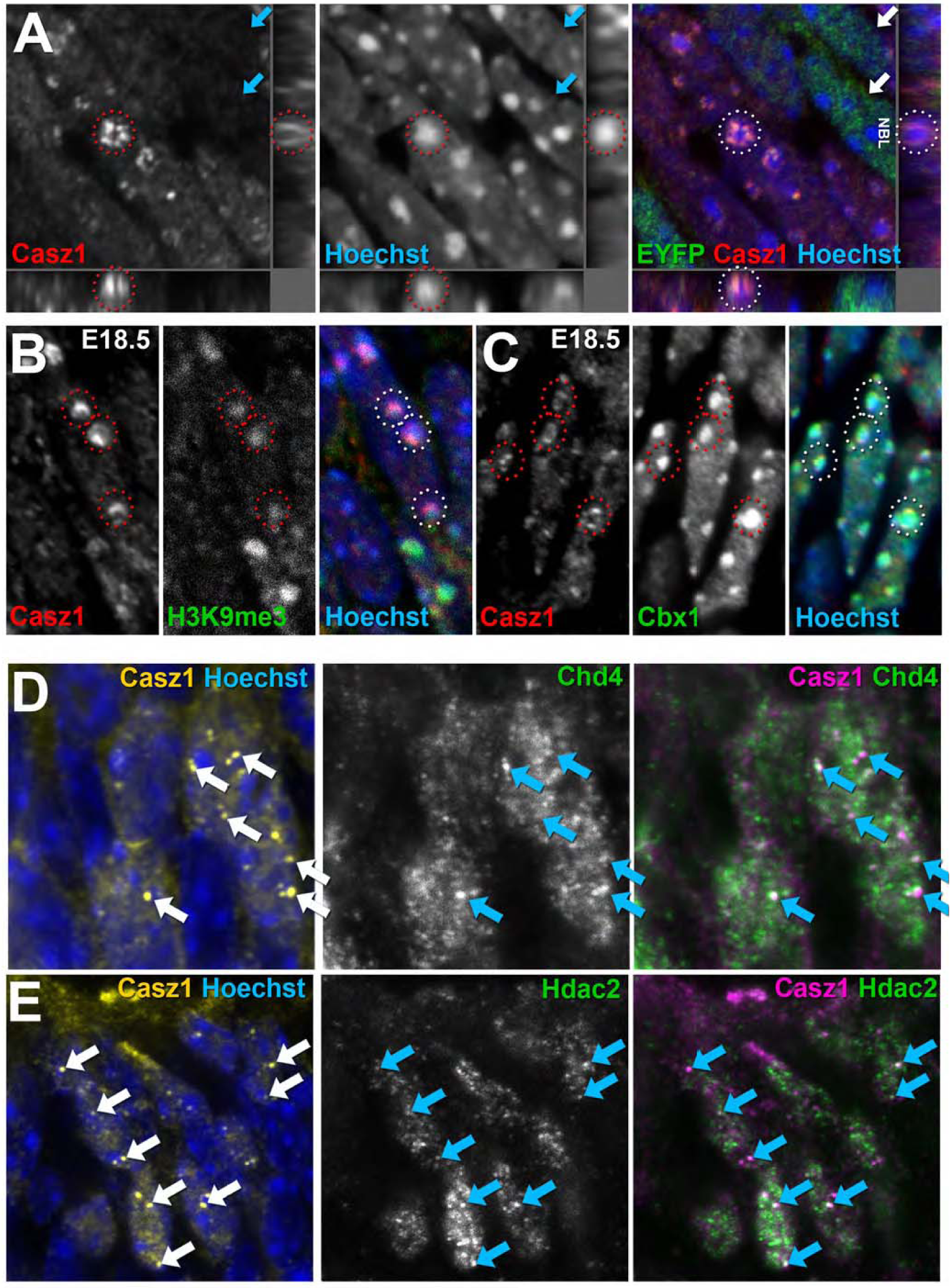
Casz1 and NuRD complex proteins localize to heterochromatin in retinal progenitors. (A) Casz1 immunohistochemistry on E17.5 *Casz1^Flox/Flox^* α*–Pax6::Cre; Rosa^YFP/YFP^* (cKO) retinal sections in mosaic regions containing recombinant (ie. EYFP+) and unrecombined (EYFP-negative) RPCs. Note that Casz1 protein staining disappears in the EYFP+ cells (arrowheads). Casz1 is localized diffusely throughout the nucleus, but is enriched on the outer margins of chromocenters (red circles). (D, E) Double-staining for Casz1 and Chd4 (D) or Hdac2 (E). Arrows mark double-positive foci.

**Fig. S2.**
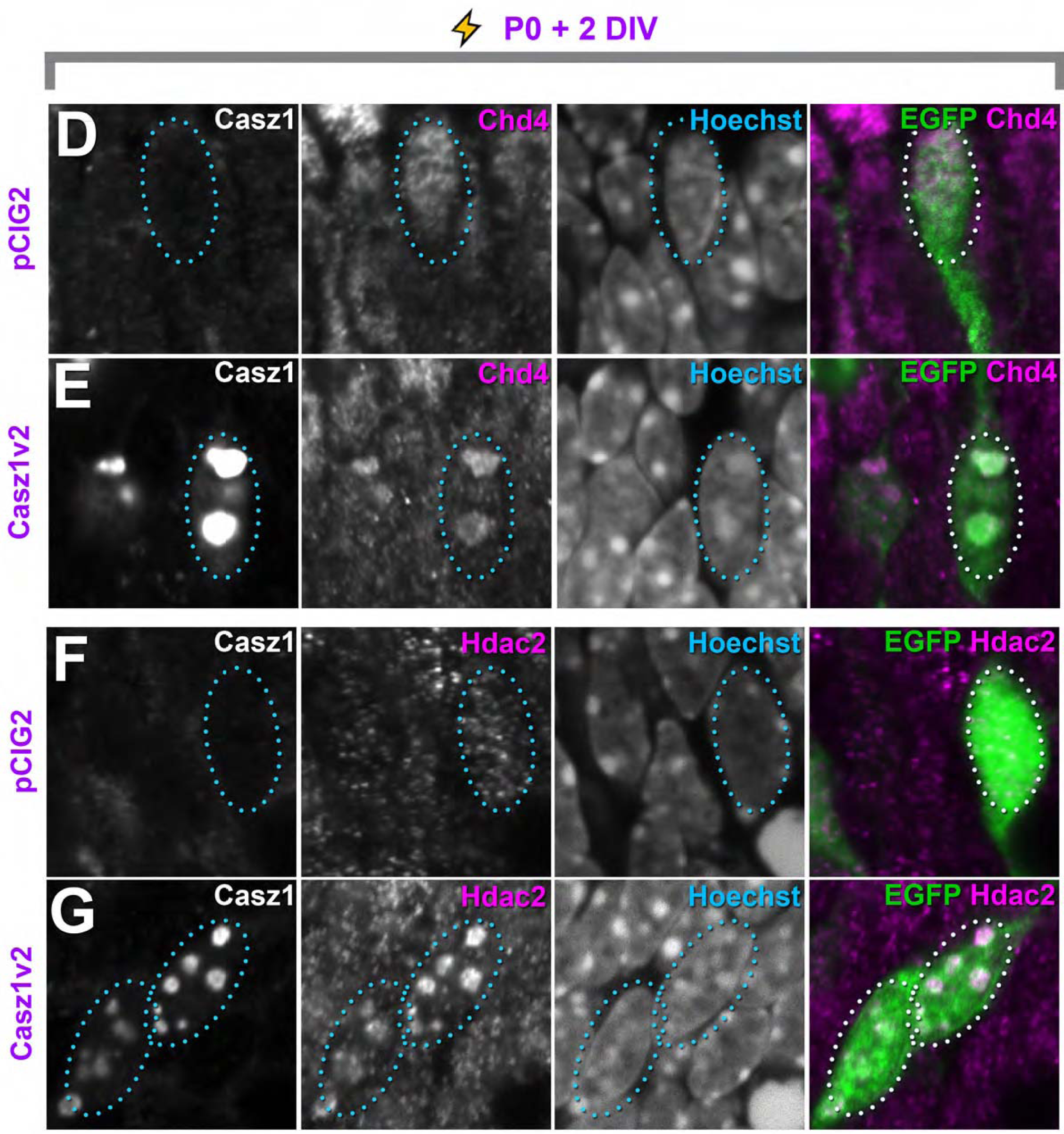
Casz1 recruits the NuRD complex to heterochromatin in retinal progenitors. (A, B) Casz1 protein localizes to the margins of chromocenters marked by H3K9me3 (A) or Cbx1 (B) in retinal progenitors. (C-F) Vector control (pCIG2; C, E) or Casz1v2 (D, F) constructs that co-express EGFP were electroporated into RPCs at P0 and cultured for 2 days in vitro. Sections were then stained for Chd4 (C, D) or Hdac2 (E, F). Casz1v2 overexpression leads to protein accumulation on chromocenters, and recruits both Chd4 and Hdac2.

**Table S1:** Statistical summary for data presented in Fig. 1D. Values refer to mean **±** SEM. NS, not significant. P-values were determined using Student’s t-test.

| Treatment | # Replicates | #Cells | % Am | % Rods | % Bi | % Mu |
| --- | --- | --- | --- | --- | --- | --- |
| *Control Flox/+* | 3 | 748 | 1.38 ± 0.99 | 79.56 ± 4.17 | 10.72 ± 5.30 | 8.34 ± 0.14 |
| *Control Flox/Flox* | 3 | 424 | 4.19 ± 1.72 | 80.08 ± 2.30 | 7.44 ± 2.47 | 7.61 ± 1.60 |
| p-value |  |  | NS | NS | NS | NS |
| *Cre Flox/+* | 3 | 621 | 4.29 ± 1.61 | 64.79 ± 0.58 | 10.57 ± 3.08 | 18.77 ± 1.45 |
| *Cre Flox/Flox* | 3 | 552 | 6.00 ± 4.40 | 49.31 ± 2.32 | 14.80 ± 1.56 | 29.40 ± 2.57 |
| p-value |  |  | NS | 0.0029 | NS | 0.0227 |

**Table S2:** Statistical summary for data presented in Fig. 3H, I. Values refer to mean **±** SEM. NS, not significant. P-values were determined using one-way ANOVA with Tukey’s post-hoc test.

| Treatment | # Replicates | #Clones | #Cells | % Am | % Rods | % Bi | % Mu |
| --- | --- | --- | --- | --- | --- | --- | --- |
| **MSCV-EGFP + vehicle** | **7** | **1128** | **1707** | **3.43 ± 0.53** | **78.00 ± 1.67** | **10.29 ± 1.74** | **8.43 ± 0.84** |
| **MSCV-EGFP + TSA** | **6** | **908** | **1503** | **3.17 ± 0.79** | **71.67 ± 0.88** | **14.50 ± 1.15** | **10.83 ± 1.08** |
| p-value vs. MSCV-EGFP veh |  |  |  | NS | NS | NS | NS |
| **MSCV-EGFP + UF010** | **6** | **1093** | **1424** | **2.30 ± 0.80** | **65.00 ± 3.60** | **7.50 ± 1.10** | **24.90 ± 3.30** |
| p-value vs. MSCV-EGFP veh |  |  |  | NS | < 0.05 | NS | < 0.0001 |
| **Cas1v2 + vehicle** | **9** | **1230** | **1736** | **0.29 ± 0.18** | **92.43 ± 1.00** | **4.14 ± 0.67** | **2.86 ± 0.26** |
| p-value vs. MSCV-EGFP veh |  |  |  | < 0.05 | < 0.01 | < 0.05 | NS |
| **Cas1v2 + TSA** | **6** | **914** | **1348** | **1.33 ± 0.61** | **75.00 ± 1.93** | **10.66 ± 1.82** | **13.17 ± 0.79** |
| p-value vs. Cas1v2 veh |  |  |  | NS | < 0.01 | < 0.05 | < 0.05 |
| **Cas1v2 + UF010** | **8** | **369** | **622** | **2.00 ± 0.89** | **62.50 ± 3.80** | **7.17 ± 0.75** | **28.67 ± 2.82** |
| p-value vs. Cas1v2 veh |  |  |  | NS | < 0.0001 | NS | < 0.0001 |

**Table S3:**
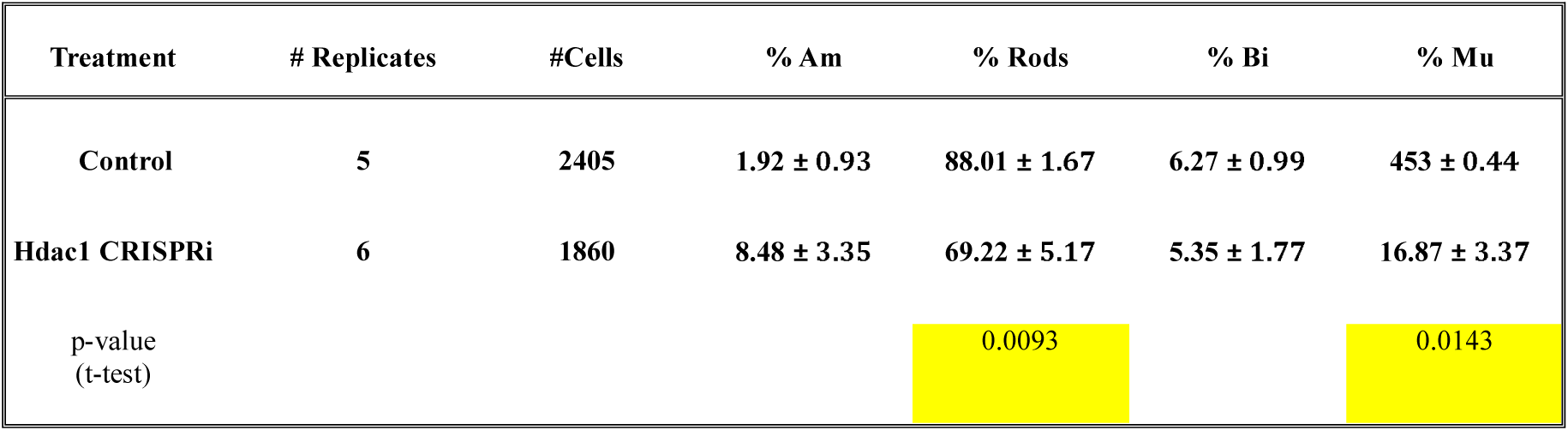
Statistical summary for data presented in Fig. 3M. Values refer to mean **±** SEM. P-values were determined using Student’s t-test.

| Treatment | # Replicates | #Cells | % Am | % Rods | % Bi | % Mu |
| --- | --- | --- | --- | --- | --- | --- |
| Control | 5 | 2405 | 1.92 ± 0.93 | 88.01 ± 1.67 | 6.27 ± 0.99 | 453 ± 0.44 |
| Hdac1 CRISPRi | 6 | 1860 | 8.48 ± 3.35 | 69.22 ± 5.17 | 5.35 ± 1.77 | 16.87 ± 3.37 |
| p-value<br>(t-test) |  |  |  | 0.0093 |  | 0.0143 |

**Table S4:** Statistical summary for data presented in Fig. 4C, D. Values refer to mean **±** SEM. P-values were determined using one-way ANOVA with Dunnett’s post-hoc test.

| Treatment | # Replicates | #Clones | #Cells | % Am | % Rods | % Bi | % Mu |
| --- | --- | --- | --- | --- | --- | --- | --- |
| **Control (pSiren)** | **3** | **495** | **757** | **1.00 ± 0** | **90 ± 1.00** | **3.33 ± 0.33** | **5.33 ± 0.67** |
| **shRing1a** | **4** | **800** | **1082** | **0.75 ± 0.25** | **79.50 ± 1.84** | **1.25 ± 0.25** | **19.00 ± 2.27** |
| p-value vs. control |  |  |  |  | 0.0172 |  | 0.0006 |
| **shRnf2-176** | **4** | **754** | **1067** | **0.50 ± 0.29** | **83.50 ± 1.32** | **2.50 ± 0.50** | **13.00 ± 0.71** |
| p-value vs. control |  |  |  |  | 0.0002 |  | 0.0171 |

**Table S5:**
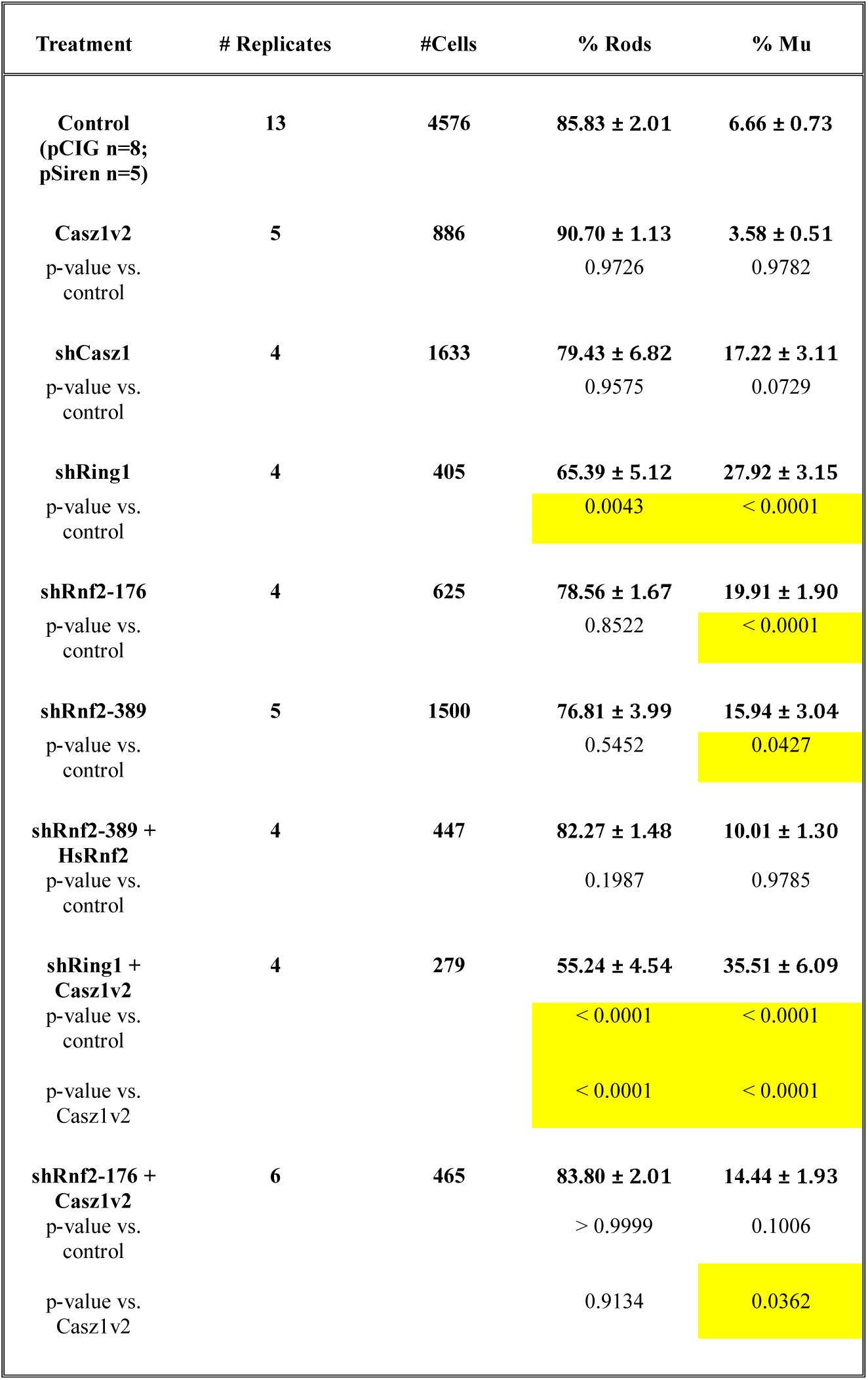
Statistical summary for data presented in Fig. 4E, F. Values refer to mean **±** SEM. P-values were determined using one-way ANOVA with Tukey’s post-hoc test.

**Table S6:**
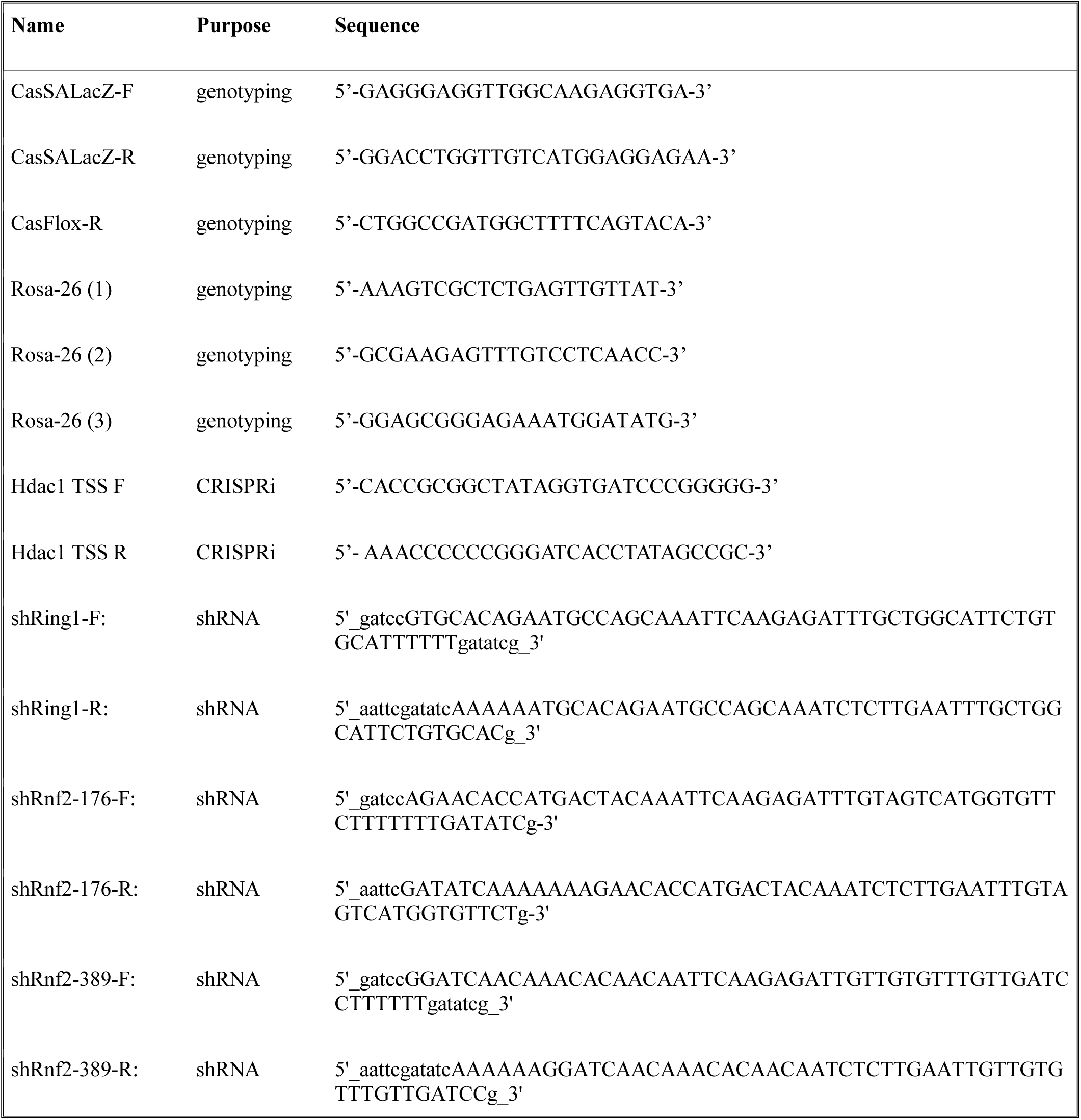
Oligonucleotide sequences.

**Table S7:** Primary antibodies and dilutions.

| Antigen | Species | Dilution | Supplier | Application |
| --- | --- | --- | --- | --- |
| Casz1 | Rabbit | 1:1000 | Seth Blackshaw Lab<br>C5-2 | IP/Western |
| Casz1 | Guinea pig | 1:200 | Johan Ericsson Lab<br>Cst95 | IHC |
| Cbx1 | Rat | 1:500 | Abcam ab10811 | IHC |
| Ccnd3 | Mouse | 1:100 | Santa Cruz D-7 | IHC |
| Chd4 | Rabbit | 1:1000 | Abcam ab72418 | IP/Western |
| Chd4 | Rabbit | 1:200 | Active Motif 39289 | IHC |
| Flag | Rabbit | 1:1000 | Cell Signaling 2368P | Western |
| GFP | Rabbit | 1:1000 | Molecular Probes<br>A11122 | IP/IHC |
| GFP | Mouse | 1:1000 | DSHB GFP-G1 | IHC |
| H2AK119ub1 | Rabbit | 1:1000 | Cell Signaling 8240 | Western |
| H3K9me3 | H3K9me3 | 1:200 | Diagenode<br>C15410056 | IHC |
| Hdac1 | Rabbit | 1:200<br>1:1000 | Bethyl A300-713 | IHC<br>Western |
| Hdac2 | Rabbit | 1:200<br>1:1000 | Bethyl A300-705 | IHC<br>Western |
| Mbd3 | Rabbit | 1:1000 | Bethyl A302-529 | Western |
| Rbbp4 | Rabbit | 1:1000 | Bethyl A301-206 | Western |
| Rnf2 | Mouse | 1:1000 | MBL D139-3 | Western |
| Sox2 | Goat | 1:500 | R&D Systems<br>AF2018 | IHC |
| Vsx2 | Sheep | 1:200 | Exalpha Biologicals<br>X1180P | IHC |

